# Effects of a complex mixture of persistent organic pollutants (POPs) on steroidogenesis in H295R cells under 10 μM forskolin stimulation - results from a pilot study

**DOI:** 10.1101/426791

**Authors:** Kareem Eldin Mohammed Ahmed, Håvard G Frøysa, Odd André Karlsen, Karin Elisabeth Zimmer, Hanne Friis Berntsen, Steven Verhaegen, Erik Ropstad, Ralf Kellmann, Anders Goksøyr

## Abstract

This study describes the utilization of an LC-MS/MS based H295R assay to assess an environmentally relevant mixture of persistent organic pollutants (POPs). H295R cells were exposed to the POP mixture in two conditions stimulated with 10 μM forskolin and unstimulated. Most importantly, the unstimulated cells responded to the low concentration of the mixture with a significant down-regulation of dehydroepiandrosterone (DHEA). This response was not observed in forskolin-stimulated cells. In stimulated H295R cells, exposure to the highest concentration showed a trend towards induced production of mineralocorticoids and glucocorticoids, although this was not significant. On the other hand, in the same exposure concentration and condition, estrogen and androgen production tended to be down-regulated. In addition to these patterns of responses being different in the stimulated vs unstimulated condition, four steroids were not detectable in the unstimulated condition.

## Introduction

The H295R cell line is considered a valuable system for the study of steroidogenic pathways, but also for the assessment of endocrine disruption caused by xenobiotics such as persistent organic pollutants or POPs (OECD, 2011; Rijk et al., 2012; Wang et al., 2014). As part of the development of a new LC-MS/MS method for studying steroidogenesis in H295R cells (Ahmed et al., 2018), we performed a pilot study where unstimulated (DMSO) H295R cells and cells stimulated with 10 μΜ forskolin were exposed to different concentrations of a total mixture of POPs, containing 29 different fluorinated, chlorinated, and brominated compounds in relative concentrations similar to those found in human blood samples (Berntsen et al., 2017). The aim was to investigate whether the stimulated cells responded to POP exposure in the same way as unstimulated cells. Although we later decided to use 1.5 μΜ forskolin as the standard condition for stimulation during POP exposure (Mohammed Ahmed et al., 2018), these data may provide an interesting background to investigate whether the higher forskolin stimulation used here influenced the response to the POP mixture in a different way than 1.5 μM forskolin.

## Materials and methods

### Chemicals

All PBDEs, PCBs and other organochlorines were originally purchased from Chiron As (Trondheim, Norway). All perfluorinated compounds (F) were obtained from Sigma-Aldrich (St. Louis, MO, USA) except perfluorohexanesulfonic acid (PFHxS) which was from Santa Cruz (Dallas, US). Hexabromocyclododecane (HBCD), phosphate buffered saline (PBS) and dimethyl sulfoxide (DMSO) were obtained from Sigma–Aldrich (Dorset, UK). Cell culture reagents were supplied by Life Technologies (Paisley, UK) and Sigma-Aldrich.

### POPs mixture

The composition of the POP mixture used in this study is presented in Table 1.

**Table 1:** Composition of the complex POP mixture used containing perfluorinated, brominated, and chlorinated compounds. Concentration are given as present in stock solution before dilution to cells.

| Perfluorinated compounds (F) | Concentration (mg/ml) | Chlorinated compounds (C) | Concentration (mg/ml) |
| --- | --- | --- | --- |
| PFOA | 1.743 | PCB 138 | 0.155 |
| PFOS | 22.348 | PCB 153 | 0.252 |
| PFDA | 0.193 | PCB 101 | 0.008 |
| PFNA | 0.507 | PCB 180 | 0.134 |
| PFHxS | 3.422 | PCB 52 | 0.006 |
| PFUnDA | 0.190 | PCB 28 | 0.008 |
| Brominated compounds (B) |  | PCB 118 | 0.045 |
| BDE-209 | 0.009 | *p,p'*-DDE | 0.339 |
| BDE-47 | 0.009 | HCB | 0.065 |
| BDE-99 | 0.004 | α-chlordane | 0.010 |
| BDE-100 | 0.002 | oxychlordane | 0.014 |
| BDE-153 | 0.001 | *trans*-nonachlor | 0.044 |
| BDE-154 | 0.002 | α-HCH | 0.005 |
| HBCD | 0.035 | β-HCH | 0.022 |
| | | γ-HCH (Lindane) | 0.005 |
|  |  | Dieldrin | 0.021 |

### Cell culture

The H295R cell line was purchased from American Type Culture Collection (ATCC). Cells were cultured in 75 cm^2^ flasks in Dulbecco’s modified Eagle medium/HamF12 (DMEM/F12) containing HEPES buffer, 1-glutamine and pyridoxine HCl (Gibco, Invitrogen, Paisley, UK). Additional supplements were added to the medium which included 1% ITS + premix (BD Biosciences, Bedford, MA) and charcoal stripped fetal bovine serum 5% (F7524, Sigma Aldrich). H295R cells were incubated at 37 °C with 5% CO_2_ in a humidified atmosphere. The medium was changed every 2–3 days and cells passaged at approximately 80% confluence by a brief exposure to 0.25% trypsin/0.53 mM EDTA (Gibco, Invitrogen). The cells were used between passages 4–6.

### Mixture exposures

After seeding H295R cells for 24 hours in 6 well plates at a cell density of 1.2 × 10^6^ cell per well, fresh medium was added containing different mixture concentrations based on tenfold dilution. Concentrations were designated (Low, Medium, High, Very High) corresponding to 1,10,100 and 1000 times the estimated concentrations in human blood (Berntsen et al., 2017). Mixtures were added to H295R cells for 48 hours. Each concentration had three biological replicates, which each was tested in three technical replicate wells.

### Cell viability

Cell viability was evaluated using Alamar Blue TM assay (Invitrogen) in 96-well cell plates (VWR, USA). Approximately 50,000 cells were seeded for 24 hours before exposure for 48 hours. Each exposure was performed in triplicate. For Alamar Blue assessment the medium was removed and replaced with 100μl of fresh medium containing alamar blue reagent for 3 hours at 5 % CO_2_ at 34 °C. A PerkinElmer (EnSpire 2300 Multilabel Reader) spectrophotometer was used to read the plates. The absorbance was read at 570nm and 600nm and viability was expressed as percentage of control (medium with 0.25 % DMSO). Triton X-100 was used as a positive control.

### LC-MS/MS

LC-MS/MS analysis was carried out on a Waters Xevo TQ-S triple quadrupole mass spectrometer that was coupled to a Waters i-class Acquity UPLC. Ionisation was achieved by electrospray ionisation (ESI) in positive and negative mode. The following LC conditions were used: chromatographic separation was achieved on a Waters Acquity BEH-C18 column (2.1 × 100 mm, 1.7 μm particle size, pore-size 130 Å). The column temperature was set at 60° C. Mobile phase A consisted of Milli-Q water with 0.05 % (vol/vol) ammonium hydroxide solution (25%), and mobile phase B consisted of methanol with 0.5 % (vol/vol) ammonium hydroxide solution (25%). The sample injection volume was 4 μl. Steroid hormones were detected and quantitated by isotope-dilution mass spectrometry by multi-reaction monitoring (MRM), as described in Ahmed et al. (2018).

## Results

Medium was collected after 48 hours exposure of H295R cells to the POP mixture for both forskolin-stimulated and unstimulated cells. We obtained LC-MS/MS data for 17 metabolites from the forskolin-stimulated H295R cells. However, in case of the unstimulated H295R cells only 13 steroids could be detected by the LC-MS/MS machine (Figure 1, and 4).

### Cell viability test

Cell viability was evaluated using the Alamar Blue assay. Fluorescence from the formed resorufin was measured and no deviation from solvent control was observed with the mixture in any concentrations, whereas the positive control Triton X-100 showed an expected decrease cell viability (Figure 3).

### Steroidogenesis

Four steroids that were not detected in the unstimulated H295R cells, appeared at detectable levels after forskolin stimulation. These were corticosterone, aldosterone, estradiol and estriol (Figure 1). In unstimulated cells exposed to the POP mixture a down-regulation of dehydroepiandrosterone (DHEA) was observed (P≤0.01) in the low concentration (Figure 1, Table 2). Otherwise, the data showed no statistically significant deviation from control.

**Table 2:** The p-values for the tests of significant effects. Each experiment is tested for a significant effect using a t-test, and the resulting p-values are adjusted for multiple testing as described in the methods section. The significance indicators of Figure 2 use the values of these tables. The coloring of the steroid names is: purple for mineral-corticoids, yellow for glucocorticoids, grey for estrogens and brown for androgens.

| Condition | Level | Pregnenolone | Progesterone | Corticosterone | Aldosterone | 17-hydroxy-progesterone | Cortisol | Cortisone | Estrone | EstroneS | Estradiol | Estriol | DHEA | DHEAS | Androstenedione | Testosterone |
| --- | --- | --- | --- | --- | --- | --- | --- | --- | --- | --- | --- | --- | --- | --- | --- | --- |
| DMSO | L | 1 | 1 | | | 1 | 1 | 1 | 1 | 1 | | | $5 \cdot 10^{-3}$ | 1 | 0.91 | 0.87 |
| DMSO | M | 1 | 1 |  |  | 1 | 1 | 1 | 1 | 1 |  |  | 0.58 | 1 | 1 | 1 |
| DMSO | H | 1 | 1 |  |  | 1 | 1 | 1 | 1 | 1 |  |  | 1 | 1 | 1 | 1 |
| DMSO | VH | 1 | 1 |  |  | 1 | 1 | 1 | 1 | 1 |  |  | 1 | 1 | 1 | 1 |
| Forskolin | L | 1 | 1 | 1 | 1 | 1 | 1 | 1 | 1 | 1 | 1 | 1 | 1 | 1 | 1 | 1 |
| Forskolin | M | 1 | 1 | 1 | 1 | 1 | 1 | 1 | 1 | 1 | 1 | 1 | 1 | 1 | 1 | 1 |
| Forskolin | H | 1 | 1 | 1 | 1 | 1 | 1 | 1 | 1 | 1 | 1 | 1 | 1 | 1 | 1 | 1 |
| Forskolin | VH | 1 | 1 | 1 | 1 | 1 | 1 | 1 | 1 | 1 | 1 | 1 | 1 | 1 | 0.48 | 1 |

Forskolin-stimulated H295R cells exposed to the highest concentration of the POP mixture showed up-regulation trends in mineral and glucocorticoids, but not statistically significant changes (Figure 1, and 2).

**Figure 1:**
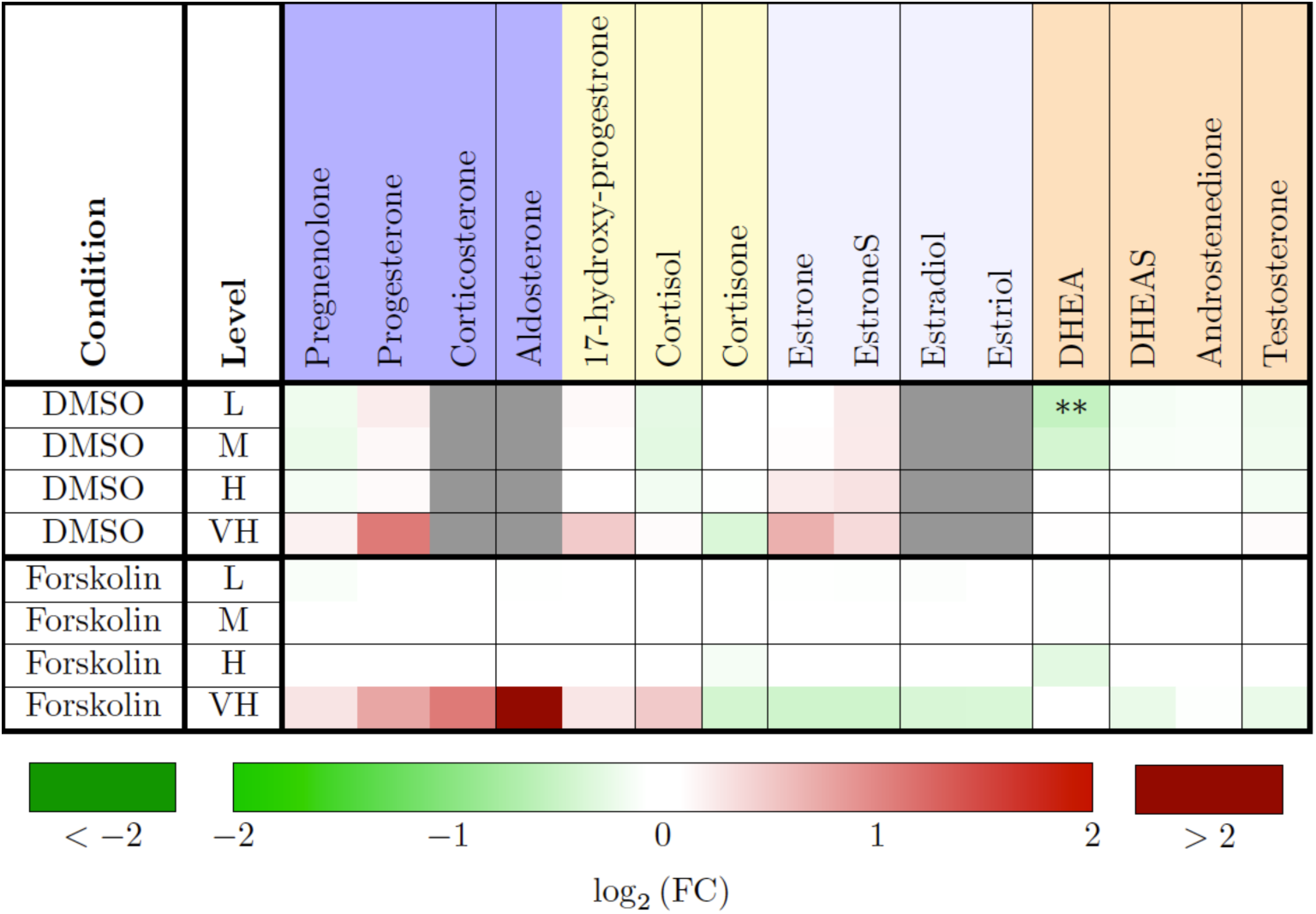
Heat map of steroid production after mixture exposure of H295R cells, with and without forskolin stimulation, indicating the fold change (FC) values for all the steroids and experiments.. The level abbreviations are L for low, M for medium, H for high and VH for very high mixture concentrations. The coloring of the steroid names is according to the groups in steroidogenesis. Each value is colored according to its log2-value where log2(FC)=0 corresponds to no effect. The grey boxes are missing data (not detected). An asterik (*) indicates a significant effect with p≤0.05 and a double asterik (**) indicates a strong significant effect with p≤0.01.

**Figure 2:**
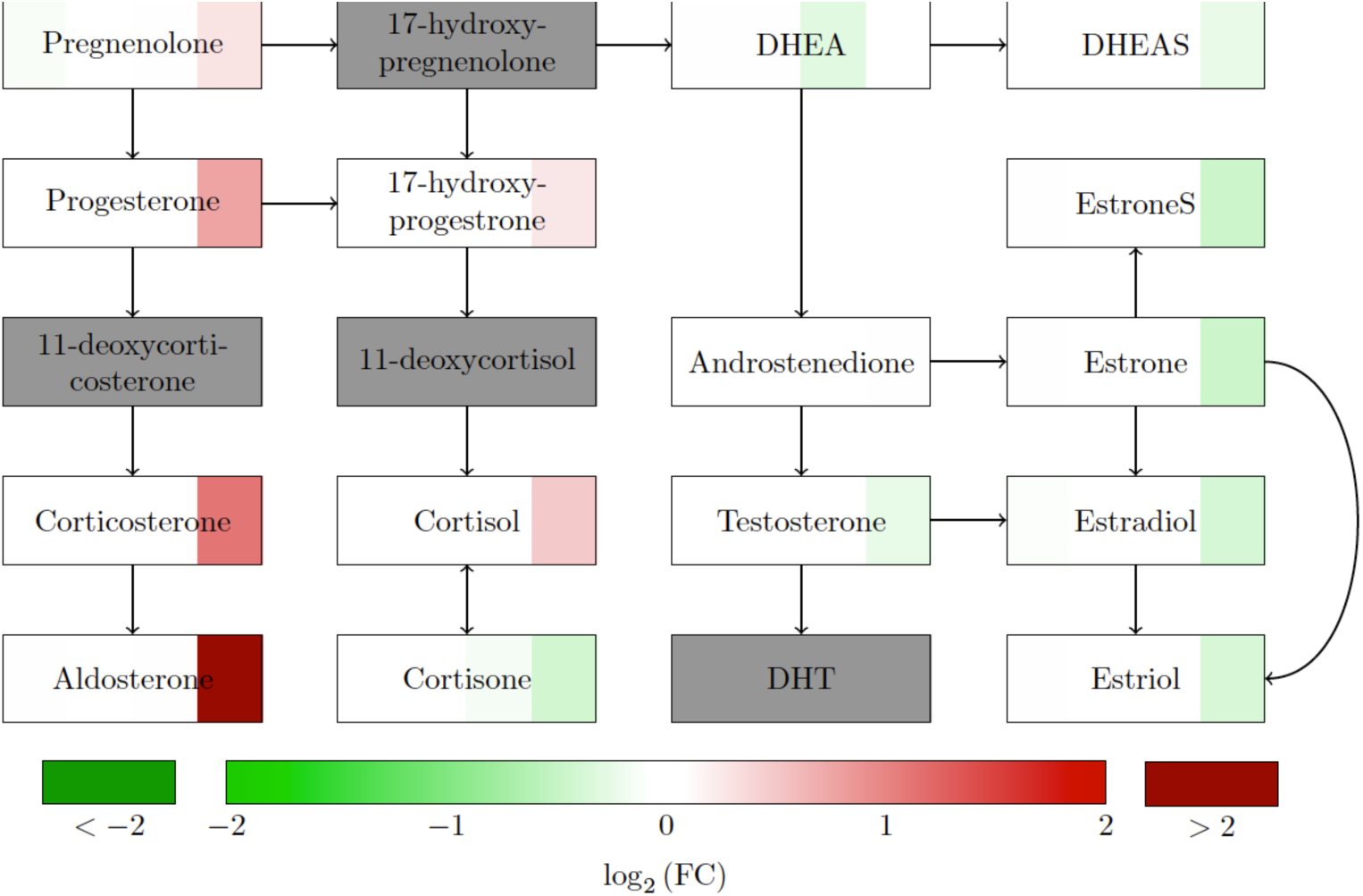
The fold change (FC) values for all the steroids in one experiment plotted with a network representation of steroidogenesis in H295R 10μM forskolin stimulated cells. The box for each steroid is divided in four parts, corresponding to low, medium, high and very high mixture concentration from left to right. Each value is colored according to its log_2_-value where log_2_(FC)=0 corresponds to no effect. The grey boxes are missing data (not detected).

## Discussion

Data from H295R cells stimulated with 1.5 μΜ forskolin exposed to the same POP mixture have been reported elsewhere (Mohammed Ahmed et al., 2018). Similar to the previous results, no significant deviations from the control were found in the present study (Figure 1, and 2). However, aldosterone and corticosterone at the highest mixture concentration indicated a strong up-regulation, although not statistically significant (Figure 4). The main difference from the responses observed under 1.5 μΜ forskolin stimulation reported in Mohammed Ahmed et al., (2018), is that there is a trend for up-regulation with regards to mineralocorticoids and to some extent glucocorticoids in the VH concentration of the mixture exposure, while androgens and estrogens in general are down-regulated at the same concentration. This may suggest that the mixture could target CYP17 function, which among others converts 17-OH-progesterone to DHEA the precursor of sex steroids (Figure 2). This may indicate that the 10 μΜ stimulation is less sensitive to responses that may be observed at the more optimal 1.5 μΜ forskolin concentration. Still, at the highest POP concentration, a similar elevation in aldosterone as with 1.5 μΜ forskolin was observed.

**Figure 3:**
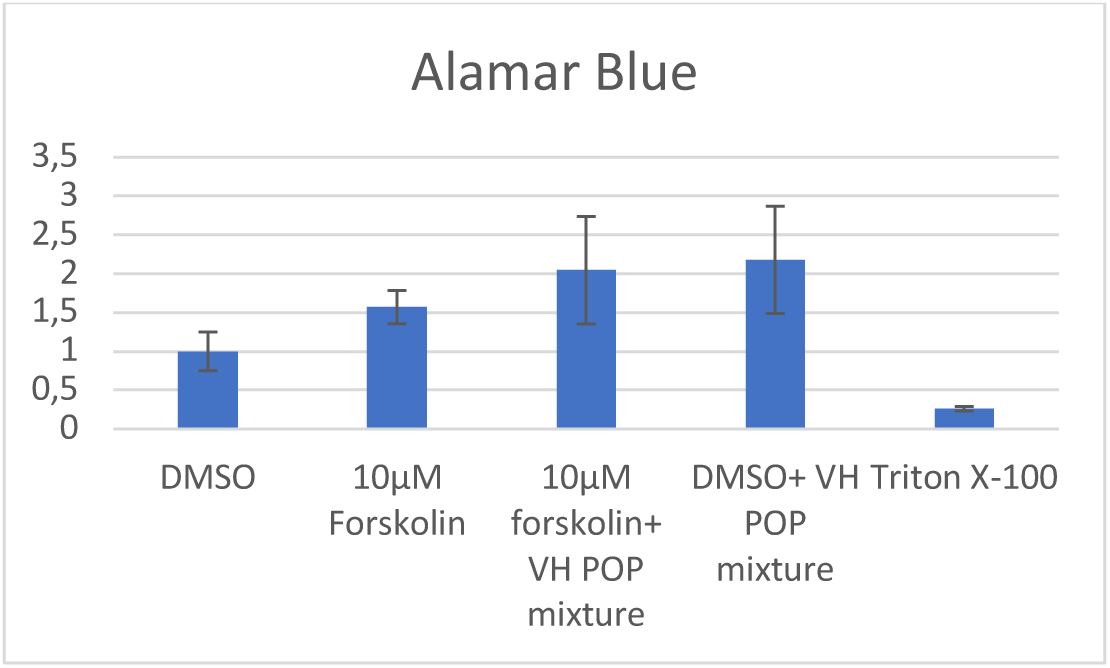
Cell viability of H295R cells after exposure to the very high concentration of POP mixtures. Cells were exposed to POPs mixture for 48 h and cytotoxicity measured Alamarblue assay. Data are expressed relative to untreated control.

**Figure 4:**
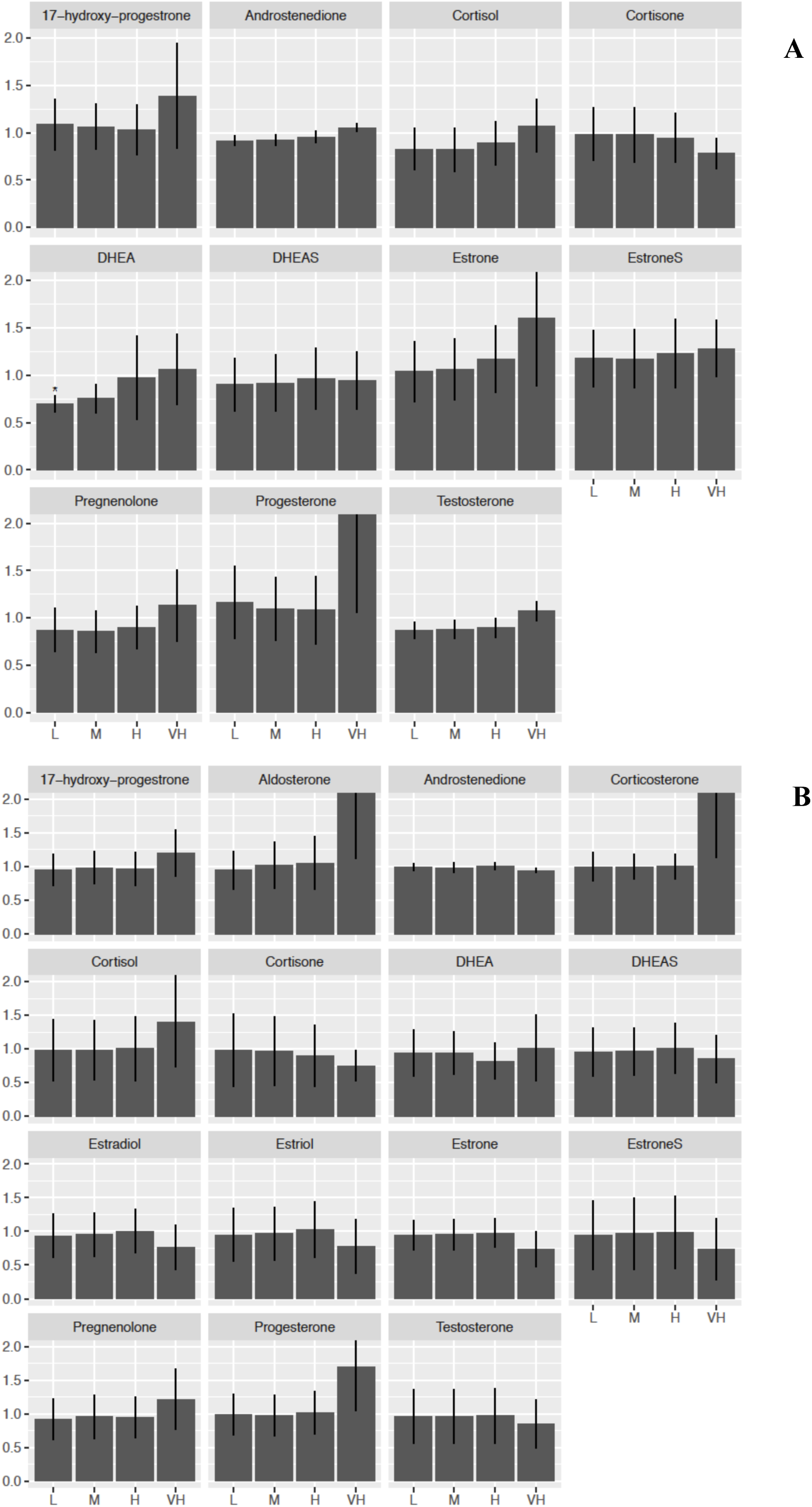
The fold change for all metabolites is plotted for A) DMSO and B) 10μΜ forskolin stimulated H295R cells exposed to the POP mixture for 48 hours. An asterisk (*) indicates a significant effect with p≤0.05. In addition, a 95% confidence interval is plotted for each value.

The data from unstimulated (DMSO treated) cells compare generally well with the results presented in Mohammed Ahmed et al., (2018), although a decrease (around 30%) in DHEA production was reported as significant here, whereas a similar decrease in Mohammed Ahmed et al., (2018) was found not to be significant.

It appears that the use of forskolin to stimulate steroidogenesis in H295R cells is necessary to utilize the full potential of the H295R assay. In the unstimulated condition several key metabolites such as corticosterone, aldosterone, estradiol and estriol are produced at undetectable levels (Figure 2; Mohammed Ahmed et al., 2018). Previously, we have found that hormone secretion in H295R cells reaches 50% saturation when cells are stimulated with 1.5 μL forskolin (Ahmed et al., 2018). This is suggested to represent an optimal condition to investigate the effect of endocrine disrupting chemical mixtures on steroidogenesis, when cells are already stimulated, but steroid production is not saturated.

Handling data with regards to H295R hormone production could be challenging as there is an observation of considerable variability among experiments conducted on different days and with different batches of cells. Hecker et al., (2006) reported a variation between experiments in testosterone production from 3.0±0.6 to 9.2±0.7 ng/ml and in estradiol form 0.21±0.05 to 0.81±0.24 ng/ml in culture medium. We have also experienced fluctuations in the levels of testosterone (53 %) and estradiol (60%) (Figure 4). This may lead to lack of power in the experimental setups, even when we normalize or combine the three experiments.

To summarize, the results from this pilot study indicate that exposing H295R cells stimulated with forskolin at 10 μM concentration to POPs induces qualitative differences in steroidogenesis compared to the unstimulated condition, and that this forskolin concentration may make the cells less sensitive to disruption of steroidogenesis by POPs, than after stimulation with 1.5 μM forskolin, when steroid production is not saturated. More data are needed to confirm this observation.

## Acknowledgements

This work is partially funded by Stress-POP project (project number: 213076) and dCod 1.0 project (project no. 248840) funded by the Norwegian Research Council. The Hormone laboratory at Haukeland University Hospital, Bergen is acknowledged for providing instruments and materials for this work. Also, Roger Lille-Langøy (staff engineer) is acknowledged for his helpful discussions.

## References

Ahmed, K.E.M., Frøysa, H.G., Karlsen, O.A., Sagen, J. V., Mellgren, G., Verhaegen, S., Ropstad, E., Goksøyr, A., Kellmann, R., 2018. LC-MS/MS based profiling and dynamic modelling of the steroidogenesis pathway in adrenocarcinoma H295R cells. Toxicol. Vitr. 52, 332–341. doi:10.1016/j.tiv.2018.07.002

Berntsen, H.F., Berg, V., Thomsen, C., Ropstad, E., Zimmer, K.E., 2017. The design of an environmentally relevant mixture of persistent organic pollutants for use in in vivo and in vitro studies. J. Toxicol. Environ. Heal. - Part A Curr. Issues 80, 1002–1016. doi:10.1080/15287394.2017.1354439

Hecker, M., Newsted, J.L., Murphy, M.B., Higley, E.B., Jones, P.D., Wu, R., Giesy, J.P., 2006. Human adrenocarcinoma (H295R) cells for rapid in vitro determination of effects on steroidogenesis: Hormone production. Toxicol. Appl. Pharmacol. 217, 114–124. doi:10.1016/j.taap.2006.07.007

Mohammed Ahmed, K.E., Frøysa, H.G., Karlsen, O.A., Blaser, N., Elisabeth, K., Berntsen, H.F., Verhaegen, S., Ropstad, E., Kellmann, R., Goksøyr, A., 2018. Effects of defined mixtures of POPs and endocrine disruptors on the steroid metabolome of the human H295R adrenocortical cell line. Manuscript submitted for publication.

OECD, 2011. Test No. 456: H295R Steroidogenesis Assay, OECD Guidelines for the Testing of Chemicals, Section 4. doi: 10.1787/9789264122642-en

Rijk, J.C.W., Peijnenburg, A.A.C.M., Blokland, M.H., Lommen, A., Hoogenboom, R.L.A.P., Bovee, T.F.H., 2012. Screening for ModulatoryEffects on SteroidogenesisUsing the Human H295R Adrenocortical Cell Line: A Metabolomics Approach. Chem. Res. Toxicol. 25, 1720–1731.

Wang, S., Rijk, J.C.W., Besselink, H.T., Houtman R., Peijnenburg, A.A.C.M., Brouwer, A., Rietjens, I.M.C.M., Bovee, T.F.H., 2014. Extending an in vitro panel for estrogenicity testing: The added value of bioassays for measuring antiandrogenic activities and effects on steroidogenesis. Toxicol. Sci. 141, 78–89. doi: 10.1093/toxsci/kfu103

